# Hydrodynamic cavitation as an energy efficient process to increase biochar surface area and porosity: a case study

**DOI:** 10.1101/280685

**Authors:** Lorenzo Albanese, Silvia Baronti, Francesca Liguori, Francesco Meneguzzo, Pierluigi Barbaro, Francesco Primo Vaccari

## Abstract

The effectivity of biochar as soil amendment is depending by its physical and chemical characteristics that are related to the type and the features of the thermal production process, such as peak temperature, heating rate, holding time, as well as from the used feedstocks. The textural characteristics of biochar in term of surface area, pore size and pore volume distribution, important for the physicochemical properties of the material, are critically dependent on the production process and the feedstock type. In this study, based on a single biochar type and a single experiment, for the first time controlled hydrodynamic cavitation was proven as a fast and effective way to enhance the biochar surface area by as much as 120%, while preserving or improving the respective chemical composition, showing far higher efficiency than the conventional increase of the peak pyrolysis temperature.

Abbreviations

AS: almond shell
BET: Brunauer–Emmett–Teller method
BJH: Barret, Joyner, and Halenda method
HC: hydrodynamic cavitation
OTP: olive-tree pruning
OS: olive stone
PW: pine wood
TP: slow thermal pyrolysis

## 1. Introduction

Since 2007, when Lehmann proposed the use of biochar as soil amendant and tool to offset the atmospheric carbon dioxide emissions (Lehmann, 2007), the international scientific community has greatly advanced in the knowledge on this topic. Many reviews state that biochar is an effective soil amendant, increases agricultural production and can effectively slow down the ongoing climate change (Cha et al., 2016; Qambrani et al., 2017).

However, the term “biochar” is still far too generic, in most of the scientific literature being defined as a result of the thermochemical decomposition in the absence of oxygen (pyrolysis) of biomass. However, the interactions between different types of process (pyrolysis or gasification), different process parameters (temperature and residence time) and different types of biomass, determine the chemical and physical characteristics of the biochar and therefore the functionality of the biochar when it is used as soil amendant (Maienza et al., 2017).

While the elemental chemical composition of the biochar is very sensitive to specific feedstock, the physical characteristics depend on the specific production process, such as slow thermal pyrolysis (TP), gasification and hydrothermal carbonization, and the main process parameters, such as working temperature, heating rate and residence time (Alburquerque et al., 2016; Cha et al., 2016; Zhang et al., 2015).

One of the most important physical parameters which determine the functionality of the biochar, is the specific surface area, usually calculated using the Brunauer–Emmett– Teller method (BET) (Brunauer et al., 1938) and expressed in square meters per unit of weight of the biochar. The higher the value, the greater the surface of the biochar that will come into contact with the soil and plants when it is used as soil amendant. Many of the exchanges and interactions between biochar, soil, plant and microorganisms occur through the surface of the biochar itself; for example, the hydrogen bonds at the surface of the biochar were proven responsible for the water absorption process (Conte et al., 2013). Most solids of high surface area are to some extent porous. The texture of such materials is hence defined by the detailed geometry of the pore space, in term of size, shape, volume (Baronti et al., 2014; Cha et al., 2016; Yuan et al., 2014).

Limiting the scope to biochar generated by means of TP processes, usually the BET surface area increases with working temperature, along with the carbon content, thus adding a greater fraction of stably sequestered carbon to the enhanced surface action. All of this, at the expense of increasing the energy consumption and decreasing the yield of biochar with regards to the input raw material (Gómez et al., 2016; Imam and Capareda, 2012; Muñoz et al., 2017).

Cavitation in liquids occurs whenever the local hydrodynamic pressure falls below the liquid’s vapor pressure at a given temperature, causing vaporization in a myriad of bubbles on the micro- to nano-scale, in turn then imploding under pressure recovery (Carpenter et al., 2017). While cavitation was once known and feared only for causing erosion damage to mechanical parts of various machinery (Dular, 2016), it can be harnessed and controlled by means of different mechanisms, such as ultrasound irradiation (Colmenares and Chatel, 2017), or pulsed laser irradiation (Ozonek, 2012; Verhaagen and Fernández Rivas, 2016). However, mechanical methods, driving controlled hydrodynamic cavitation (HC) processes, have been recognized as the most energy efficient, robust, and easily scalable. Invariably, they generate HC processes by means of relative acceleration between the liquid and mechanical parts, such as circulating the liquid across a nozzle, in turn driving pressure drop according to Bernoulli’s equation (Gogate and Pandit, 2011) When the generated vapor cavities meet a zone with higher pressure, such as downstream a nozzle, they undergo fast collapse (implosion), thereby concentrating the kinetic energy of the bulk medium into very small size hot spots. Temperature and pressure inside a collapsing bubble increase dramatically up to 5,000-10,000 K and 300 atm, respectively, due to the work done by the liquid to the shrinking bubble, producing very strong shear forces, micro-jets and pressure shockwaves (Pawar et al., 2017; Yasui et al., 2004).

Different HC regimes will be practically identified based on the values assumed by a single dimensionless parameter, *i.e.* the cavitation number (*σ*) derived from Bernoulli’s equation. It represents the ratio between the pressure drop needed to achieve vaporization, and the specific kinetic energy at the cavitation inception section (Šarc et al., 2017; Yan and Thorpe, 1990), as per Eq. (1).

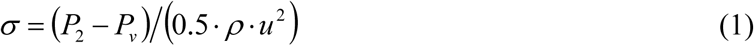

where *P*_*0*_ (Nm^−2^) is the average pressure downstream of a cavitation reactor (the recovered pressure), such as a Venturi tube or an orifice plate, where cavitation bubbles collapse, *P*_*v*_ (Nm^−2^) is the liquid vapor pressure (a function of the average temperature for any given liquid), ρ (kgm^−3^) is the liquid density, and *u* (ms^−1^) is the flow velocity through the nozzle of the cavitation reactor, the latter also depending on the pump’s inlet pressure. However, a thorough discussion of the issues raised by the use of the cavitation number as per Eq. (1) has been recently advanced (Šarc et al., 2017), along with the issuance of a comprehensive set of suggestions and recommendations, aimed at improving the understanding and repeatability of HC processes and experiments. In particular, it was shown that by changing the very definition of the different parameters, included in the expression for *σ*, could lead to differences in the respective resulting values as large as two orders of magnitude. Specific prescriptions concerned the pressure *P*_*0*_ and the velocity *u*, which should always be measured, respectively, downstream of the cavitation constriction and through it.

In order to comply with the above-mentioned recommendations, as well as to allow repeatability, the relevant details of the HC device and related sensors are supplied in Section 2.

As a rule, with Venturi tube reactors, and definition of parameters, adopted in this study, developed cavitation, with fairly strong and frequent collapses, arise within the range 0.1<*σ*<1 (Bagal and Gogate, 2014; Gogate, 2002), which is the range adopted in the experiments described in this study. Moreover, the collapse of cavitation bubbles, with inner temperatures over 2,500 K, causes the formation of powerful oxidants, such as hydroxyl radicals (·OH, oxidation potential 2.80 eV), as a result of water splitting (Ciriminna et al., 2017; Yasui et al., 2004). However, without the assistance of further advanced oxidation processes, such as due to additives like hydrogen peroxide, ozone, or Fenton reagents, the extent of oxidation of the bulk liquid medium is quite limited (Yusaf and Al-Juboori, 2014). Indeed, no oxidation was observed either in wort or in final beer produced by means of HC-assisted brewing processes (Albanese et al., 2017a).

Several different HC-driving mechanical devices can be enumerated, such as rotor-stator arrangements (Badve et al., 2015; Petkovšek et al., 2015), mechanical constrictions and nozzles, such as orifice plates (Gogate and Kabadi, 2009; Rudolf et al., 2017), Venturi tubes (Šarc et al., 2017; Zamoum and Kessal, 2015), or combination of both (Li et al., 2017). However, in the treatment of liquids containing significant amounts of solid particles, Venturi tube reactors offer minimization of the obstruction risk, simplicity and robustness (Albanese et al., 2015).

In this work, for the first time the emerging technique of controlled hydrodynamic cavitation (HC) is experimentally investigated, about its applicability to increase the surface area of the biochar, along with its performance compared to the conventional solution of increasing the peak temperature of TP processes.

## 2. Materials and methods

### 2.1 HC device and process

The experimental device (Fig. 1), implementing the HC-based process on a pilot scale, includes a closed hydraulic loop with total volume capacity around 230 L, powered by a centrifugal pump (7.5 kW nominal mechanical power, rotation speed 2900 rpm), the same designed to produce beer wort in past studies (Albanese et al., 2018, 2017a, 2017b).

**Figure 1.**
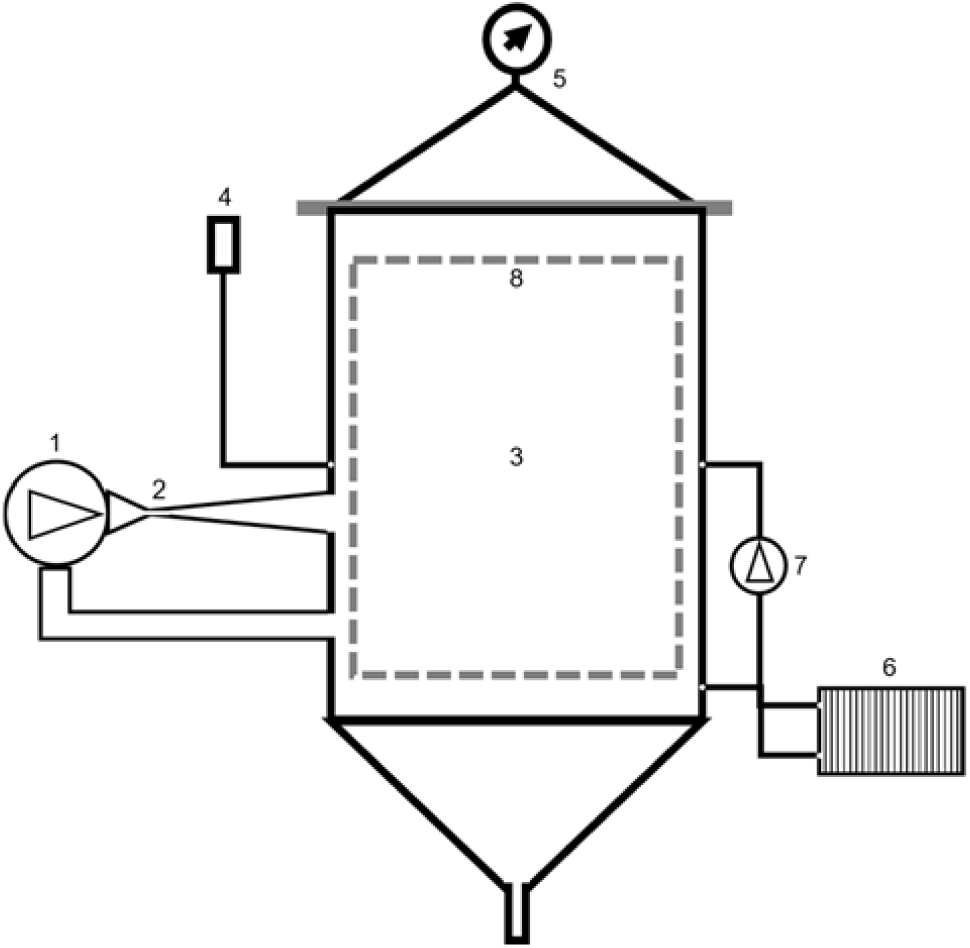
Simplified scheme of the experimental HC-based installation. 1 – Centrifugal pump, 2 – HC reactor, 3 – main vessel, 4 – pressure release valve, 5 – cover and pressure gauge, 6 – heat exchanger, 7 – circulation pump, 8 – malts caging vessel. Other components are commonly used in state-of-the-art hydraulic constructions

Any surface in contact with the circulating solid-liquid mixture was crafted in food-quality stainless steel (AISI 304), with 2 mm minimum thickness. The cavitation reactor, in the form of a circular-section Venturi tube, described in detail in a past study (Albanese et al., 2015), was preferred over an orifice plate due to the potential obstruction produced by the circulating solid particles. During the test run, the circulating solid-liquid mixture was exposed to the atmospheric pressure; however, the device was designed to allow additional hydraulic pressure by means of a tunable pressure release valve, by means of which the cavitation number *σ* can be tuned across a wide range of values through the *P*_*0*_ term in Eq. (1) (Soyama and Hoshino, 2016).

While circulating, the mixture heats up, due to the conversion of impeller’s mechanical energy into thermal energy; the heating source concentrates just downstream the cavitation reactor, where vigorous internal friction occurs due to the cavitation process.

The HC experimental unit was equipped with standard thermometer and manometer sensors. In particular, the wort pressure *P*_*0*_, contributing to the cavitation number in Eq. (1), was measured far downstream the nozzle of the Venturi reactor, as shown by the location of the pressure gauge, thus complying with the recommendations mentioned in Section 1 (Šarc et al., 2017).

The power and electricity consumed by the HC-assisted process were measured by means of a commercial three-phase digital power meter, which was also used to assess the flow velocity through the nozzle of the cavitation reactor, according to the procedure explained in a past study (Albanese et al., 2017a), as well as again complying with the aforementioned recommendations.

After sieving the original sample with a linear dimension ≤ 3 mm, a mass of 5.73 kg of biochar was obtained, such sieving being required by the use of a closed impeller pump, which could not manage greater size material. Then, the biochar was mixed with a volume of unfiltered tap water of 130 L, *i.e.* at the concentration of 4.4%, within the first 3 min of the test run.

Original water acidity was neutral, with pH 7.0±0.2, and the heating rate was unconditioned, in order to avoid additional energy consumption for cooling, resulting from the balance between the mechanical energy supplied by the pump impeller and the heat loss through the uninsulated walls of the hydraulic circuit. No additives were used for the test.

### 2.2 Biochar samples

The biochar sample used for the HC process was produced from coppiced woodlands (beech, hazel, oak and birch) through a pyrolysis process with an average residence time of 3 hours at 550°C in a transportable kiln provided by Blucomb (http://www.blucomb.com/) of 0.8 m in diameter, holding around 30 kg of feedstock. C, N, H and S contents of biochar, both original and during the HC process, were determined, after complete removal of inorganic C with acid, using an Elemental analyzer (Perkin-Elmer, model 2400). The biochar samples were screened through a 2 mm sieve and oven dried at 105°C for 24 h; the dry samples were acid digested in a microwave oven (CEM, MARSX press) according to the EPA method 3052 (USEPA, 1995). The solutions obtained after the mineralization were filtered (0.45 mm PTFE) and diluted. Biochar pH was measured in a 1:4 biochar/water suspension. Ash content was measured by heating the samples at 800 °C for 4 h. The oxygen content was determined by a mass balance.

Finally, the acidity of the water-biochar mixture was measured by means of pH-meter (Hanna Instruments, Padova, Italy, model HI 99151) with automatic pH calibration and temperature compensation.

### 2.3 Biochar textural properties

The textural properties of the biochar before and after cavitation were determined by nitrogen adsorption and desorption at −196°C using a Micromeritics ASAP 2020 volumetric sorption analyzer. Prior to gas adsorption, the samples were activated in situ by heating to 300°C, at a rate of 5°C/min under vacuum for 12h, using the degassing system of ASAP 2020.

The surface area was calculated by using the BET method and Langmuir method applied to nitrogen adsorption data in the relative pressure (P/P°) range of 0.06-0.40. The total pore volume was determined from the amount of nitrogen adsorbed at P/P°=0.98. The pore size distribution was obtained using the Barret, Joyner, and Halenda (BJH) method (Thommes et al., 2015). The assessment of microporosity was carried out by *t*-plot method. A minimum equilibrium interval of 30 s with a maximum relative tolerance of 5% of the targeted pressure and an absolute tolerance of 5 mmHg were used.

## 3. Results and Discussion

Table 1 shows the basic features of the HC processes, *i.e.* the evolution of temperature and specific energy (consumed electricity per unit mass of biochar), along with the main textural and chemical properties of the biochar throughout the process.

**Table 1.** Basic features of the HC processes, and main textural and chemical properties of the biochar throughout the process.

| Process time<br>(min) | Temp.<sup>a)</sup><br>(°C) | Spec. energy<sup>b)</sup><br>(kWh/kg) | BET (micro, meso)<sup>c)</sup><br>(m<sup>2</sup>/g) | pH<sup>d)</sup> | pH<sub>24</sub><sup>e)</sup> | C<br>(%) | H<br>(%) | H/C | O<br>(%) | O/C | N<br>(%) | Ash<br>(%) |
| --- | --- | --- | --- | --- | --- | --- | --- | --- | --- | --- | --- | --- |
| 0 | 22.0 | 0.000 | 59.95 (35.38, 24.57) | 9.00 |  | 68.60 | 1.00 | 0.01 | 29.91 | 0.44 | 0.49 | 42.49 |
| 5 | 25.5 | 0.109 | 80.25 (47.02, 33.23) | 9.35 | 9.47 | 69.74 | 1.24 | 0.02 | 28.62 | 0.41 | 0.40 | 34.01 |
| 20 | 35.5 | 0.443 | 95.55 (60.30, 35.25) | 9.47 | 9.59 | 61.05 | 1.32 | 0.02 | 37.03 | 0.61 | 0.59 | 31.36 |
| 30 | 42.0 | 0.656 | 113.32 (71.59, 41.73) | 9.47 | 9.63 | 65.65 | 1.40 | 0.02 | 32.37 | 0.49 | 0.58 | 31.21 |
| 45 | 50.5 | 0.973 | 130.76 (86.06, 44.70) | 9.55 | 9.65 | 52.26 | 1.30 | 0.02 | 45.83 | 0.88 | 0.61 | 32.16 |
| 60 | 58.5 | 1.284 | 135.27 (89.00, 46.27) | 9.49 | 9.69 | 57.87 | 1.16 | 0.02 | 40.38 | 0.70 | 0.59 | 32.28 |
| 75 | 65.5 | 1.590 | 129.21 (86.77, 42.44) | 9.77 | 9.64 | 53.50 | 1.15 | 0.02 | 44.79 | 0.84 | 0.56 | 34.05 |
| 90 | 72.5 | 1.892 | 131.05 (86.21, 44.84) | 9.76 | 9.69 | 56.09 | 1.25 | 0.02 | 42.09 | 0.75 | 0.58 | 29.82 |

a) Temperature of the water-biochar mixture.
b) Consumed electricity divided by the mass of processed biochar.
c) Surface area calculated by using the BET method, surface area of micropore and mesopore calculated by the *t*-plot method.
d) pH measured in the water-biochar mixture.
e) pH measured 24 hours after processing.

### 3.1 HC process

Due to unconditioned heating and negligible heat losses within the moderate temperature range (22 to 72.5°C), as shown in Table 1, both temperature and specific energy quantities are approximately linearly dependent on time.

While the biochar powder was oven-dried after the HC process for analytical purposes, the respective consumed energy was not accounted for, because biochar is not necessarily spread in the soil as a dry powder. Indeed, as biochar is a fine powder with pyrophoric properties, it is often quenched with water in order to avoid auto-ignition and allow ease handling prior to its end use (Peters et al., 2015). Moreover, very likely, the concentration of biochar could be increased over the level of 4.4% used in this experiment, resulting in lower specific energy consumption, which will be the subject of further research.

The cavitation number, computed as per Eq. (1), was substantially stable, between 0.073 and 0.1, throughout the process. Constant power consumption proved the regime of developed cavitation, despite the relatively low values of the cavitation number, as expected based on the abundance of condensation nuclei supplied by the biochar powder.

### 3.2 Biochar textural properties

Recalling the methods described in subsection 2.3, almost all the N_2_ adsorption-desorption isotherms for the biochar carbons before and after the HC process exhibited a type I isotherm with a low knee at low pressures, and only a few samples showed a mixture of type I and IV isotherm and of type I and II.

An typical example of N_2_ adsorption-desorption isotherm of biochar collected after HC process time was showed in Fig. 2(a), while Fig. 2(b) shows all the N_2_-adsorption isotherms. All N_2_-adsorption-desorption isotherms and the BJH Desorption dV/dLog(w) Pore Volume schemes for biochar, throughout the process and starting from the sample collected after 45 min of process time, are shown in Fig. S1-S7, available in the Supplementary material. Table S1 in the Supplementary material as well as the BET surface area other parameters such as the Langmuir surface area, the *t*-plot micropore area, the pore volume, and the average pore width are shown.

**Figure 2.**
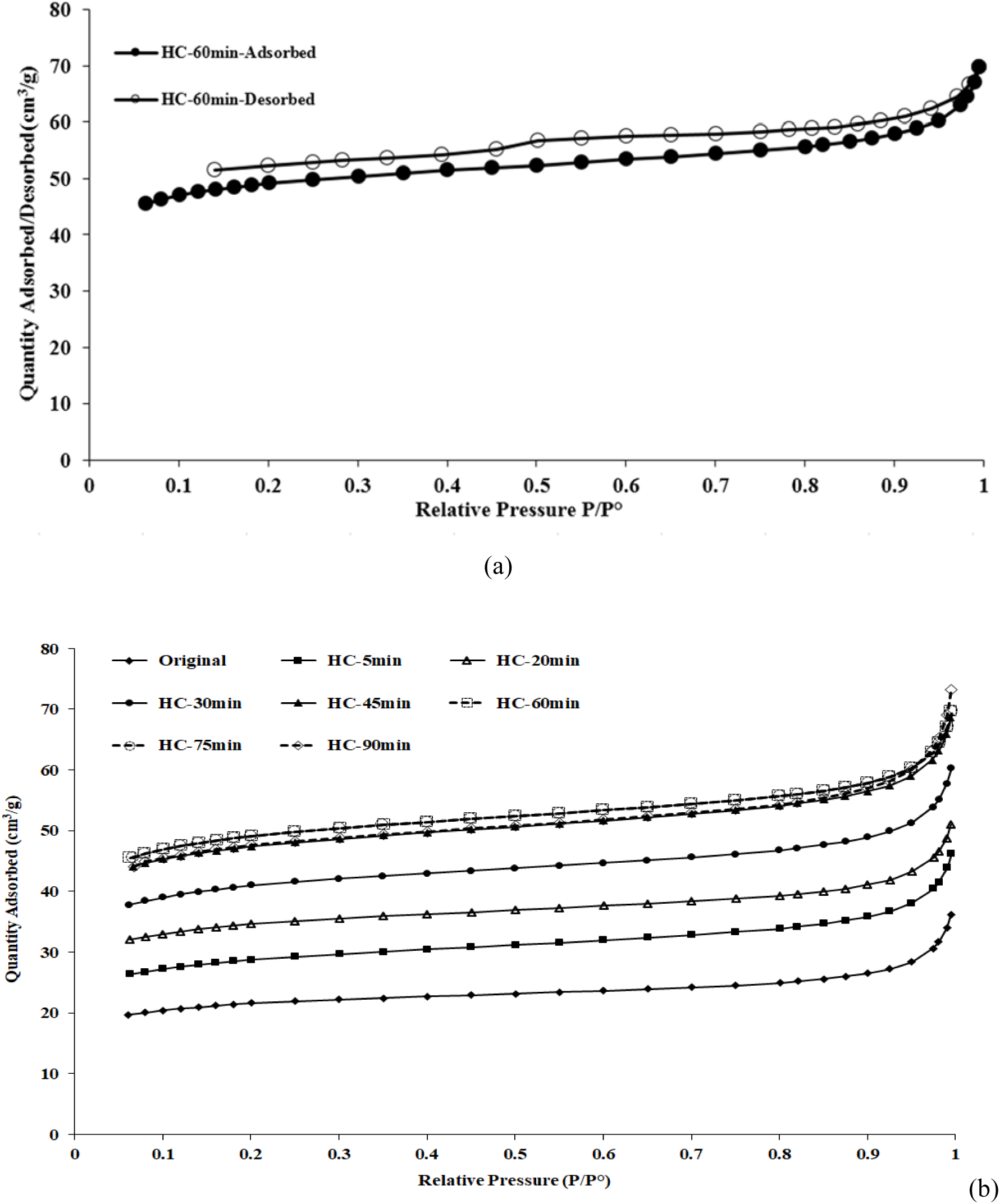
N_2_-adsorption-desorption isotherms at 77.4 K for biochar after 60 min of HC process time (a), and all the N_2_-adsorption isotherms (b).

According to the IUPAC classification (Thommes et al., 2015), type I isotherm can be associated with microporous structure having relatively small external surface, the limiting uptake of nitrogen being governed by the accessible micropore volume rather by the internal surface area. The mixture of type I and IV or II can be attributed to a material with micro and mesoporosity, with the initial part of the isotherm attributed to a monolayer-multilayer adsorption and a multilayer formation at relatively high partial pressure. Hysteresis loop exhibited the shape *H4*, often associated with narrow-slit-like pores, but in the case of the type I isotherm it is indicative of microporosity.

Those isotherms clearly showed the predominantly microporous nature of biochar elements, with some mesoporous leading the gradual increase in adsorption after the initial filling of the micropores, followed by faster enhancement near saturation. The BET surface area, as shown in Table 1 showed a very fast increase in the first 5 min of HC processing, at the rate of 6.8%/min (about 60 to 80 m^2^/g, *i.e.* +34%), later increasing about linearly with time at a reduced rate of 1.6%/min up to 45 min of process time. Later, this parameter stabilized in the narrow range 129 to 135 m^2^/g, *i.e.* around 120% over the original level, with the peak level of 135.27 m^2^/g observed after 60 min of process time.

After the first 5 min, during which the relative contributions to the BET growth from micropores and mesopores were comparable, most of the contribution, up to 45 min of process time, came from the increasing micropores surface, at the level of 77%, against only 23% from the mesopores surface. The contribution of micropores to the observed increase in surface area, is further confirmed by other data shown in Table S1 such as the Langmuir surface area, the *t*-plot micropore area, and of the *t*-plot micropore volume. The average mesopore width, calculated with the BJH method, decreased from a value of 8 nm before cavitation to a value of 6.3 nm after 5 min of process time, later stabilizing in a narrow range around 6 nm throughout the process.

Thus, the very fine cracking of the solid surface of the biochar elements arises as a relevant physical effect, in agreement with the proven ability of HC processes to assist in synthesizing nanomaterials, as a result of liquid jets and pressure shockwaves at the collapse of cavitation bubbles close to solid surfaces (Carpenter et al., 2017).

### 3.3 Biochar chemical properties

The carbon content (Table 1) oscillates around 65% during the first 30 min of process time, later decreasing abruptly by about 20%, *i.e.* down to about 52%, in the time lapse 30 to 45 min, finally stabilizing around 55%. The hydrogen content steadily increases from 1.00% to 1.40% up to 30 min of process time, later decreasing to levels in the range 1.15% to 1.25%. The oxygen content increases in two steps: the first, from 5 min to 20 min (28.62% to 37.03%), the second, from 30 min to 45 min (32.37% to 45.83%), later stabilizing around 42%. The nitrogen content decreases from 0.49% to 0.40% in the first 5 min, then grows up to an average level of 0.58% for the remainder of the process. The ash content decreases abruptly in the first 5 min of process time (42.49% to 34.01%), later stabilizing around 32%. Finally, the pH grows slowly from 9.35 after 5 min, up to 9.77 after 75 min; 24 hours after processing, pH of any sample up to the 60 min one marginally increases, but remains under the level of 9.7.

### 3.4 Comparative analysis of process yield

The objective comparative assessment of the performance of the HC process required a measure for the process yield. For this purpose, the yield of both TP and HC processes, at any of the respective stages, was computed as the BET surface area percentage change per unit specific energy consumption (consumed electricity per unit mass of biochar, thus including declining biochar yield for TP processes), multiplied by the percentage change of the carbon content. The reference levels were chosen at the lowest working temperature of TP processes, or at the beginning of the HC process.

More in detail, the yield for TP processes was computed according to Eq. (2-5):

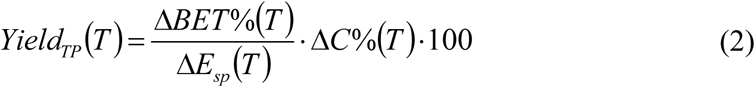

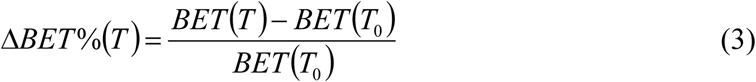

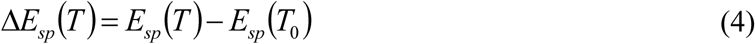

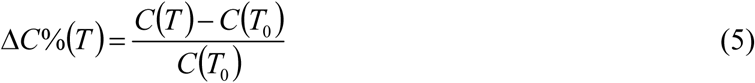

where the suffix *TP* stands for thermal pyrolysis, *T* is the working temperature (in °C) for a specific TP process, and *T*_*0*_ is the minimum working temperature, assumed as reference. *BET(T), E*_*sp*_*(T)* and *C(T)* are the BET (m^2^/g), the specific energy per unit mass of biochar (kWh/kg), and the percentage carbon content, respectively, at working temperature *T (°C)*.

The yield for the HC process was computed according to Eq. (6-9):

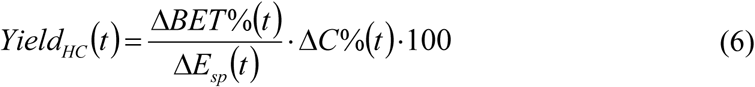

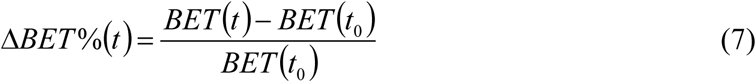

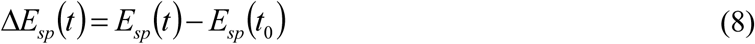

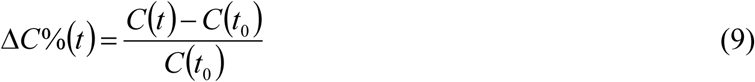

where *t* (min) is the process time from beginning, i.e. from time *t*_*0*_.

Due to the scarcity of published data including simultaneously the BET surface area, the biochar yield and carbon content, along with the energy consumption for electrically-powered pyrolysis devices, Fig. 3(a) shows results derived from data drawn from a single publication, based on pine wood (PW), olive-tree pruning (OTP), almond shell (AS) and olive stone (OS) as raw materials (Gómez et al., 2016). In particular, the energy consumption was computed adding the energy required to heat the reactor to the working temperature, *i.e.* 350, 450 and 550°C, which was assumed to be constant, irrespective of the raw material, to the energy demand for the pyrolysis process, having residence time of 15 min, at any given working temperature. Data in Fig. 3(a) are represented in correspondence of the pyrolysis temperatures of 450 and 550°C, showing the process yield in the temperature ranges 350-450°C and 450-550°C, respectively. Fig. 3(b) shows the corresponding results for the HC process, up to 45 min of process time.

**Figure 3.**
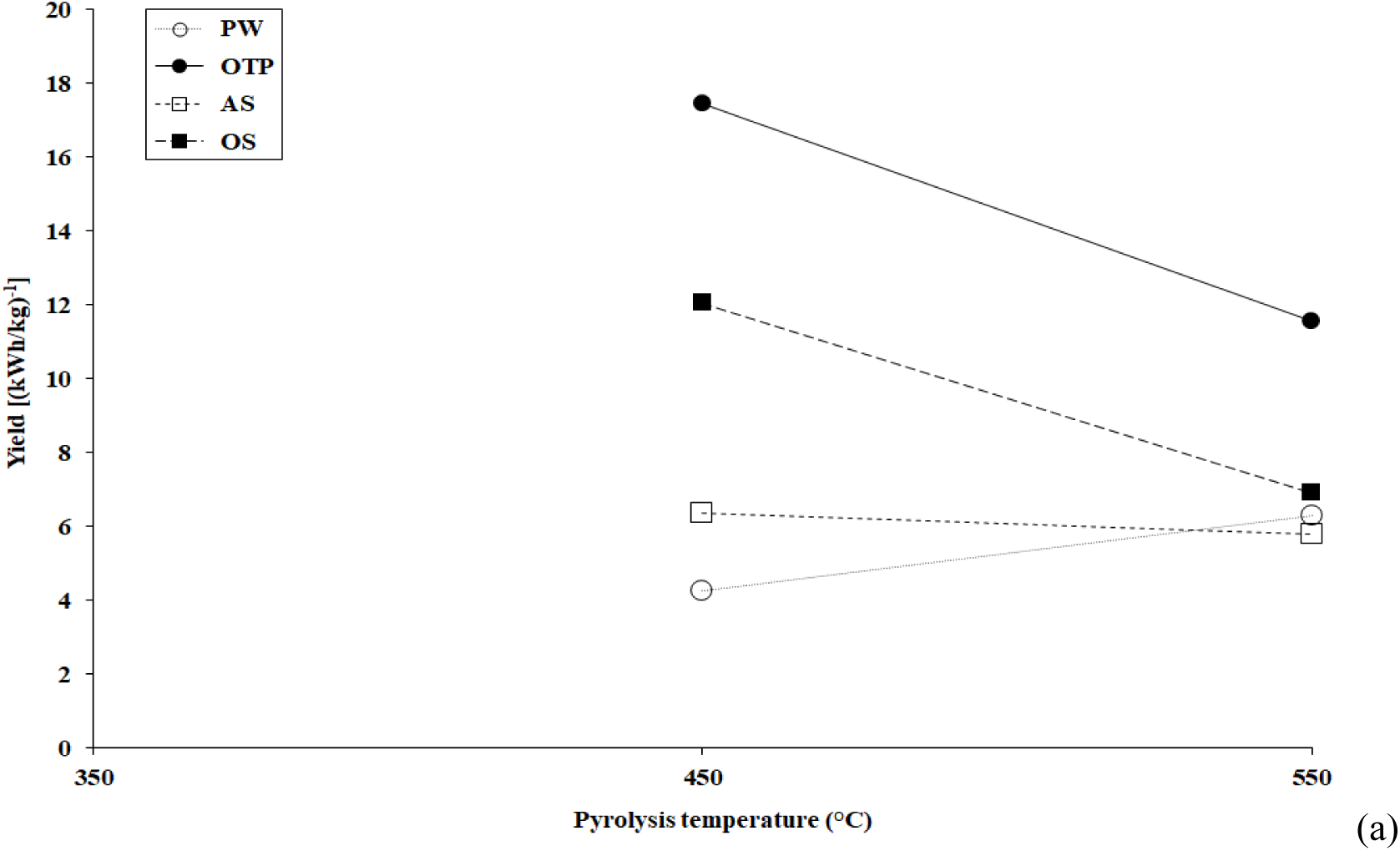
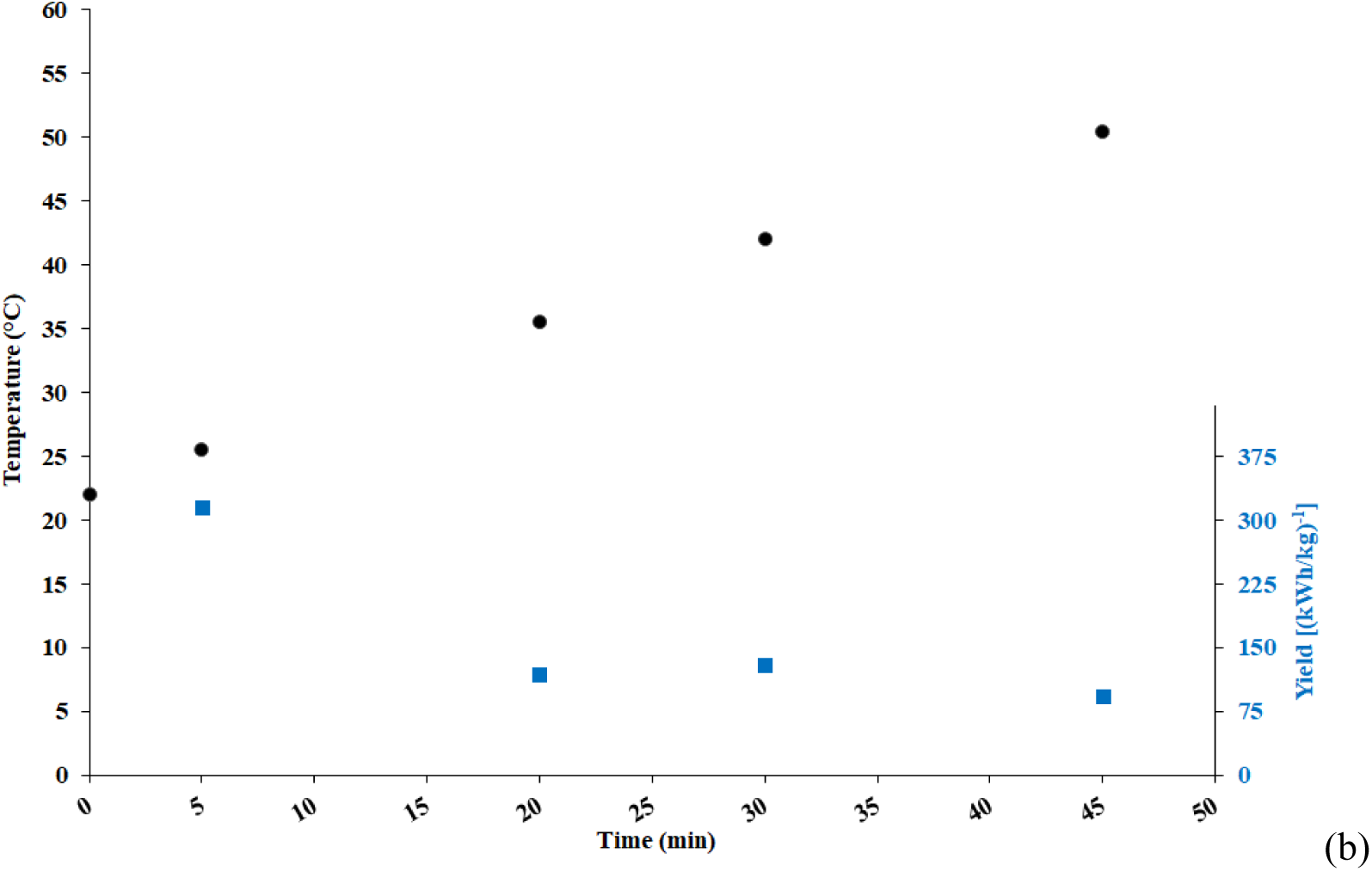
TP process yields based on literature data (Gómez et al., 2016), for PW, OTP, AS and OS as raw materials (a), and corresponding results for the HC process, up to 45 min of process time (b).

Process yields from conventional TP are lower than 18 (kWh/kg)^−1^, peaking for OTP in the temperature range 350-450°C, converging towards the narrow interval 5.9- 6.9 (kWh/kg)^−1^ in the temperature range 450-550°C for PW, AS and OS. The HC process yields change from a minimum of about 92 (kWh/kg)^−1^ for a process time of 45 min to as much as about 315 (kWh/kg)^−1^ in the first 5 min of the process. A local maximum of the process yield is located at a process time of 30 min, at the level of 130 (kWh/kg)^−1^, *i.e.* between 7 and 30 times higher than the yields assessed for the considered TP processes.

Thus, within the limit of the discussed case study, the HC process appears as an extremely energy efficient way to increase the surface area (BET), while preserving the carbon content, with wide margins for further improvement, starting from the use of a greater concentration of biochar in the HC process, as mentioned in subsection 3.1. However, further comparative performance studies are recommended, including far more data for both TP and HC processes, along with further research on the sensitivity to different definitions for process yield.

### 3.5 Optimization of the HC process

Among the generally accepted quality criteria that biochar should fulfill for carbon sequestration and agricultural uses, the following can be verified against the available data: C content >50%, H/C <0.7, O/C <0.4, and BET >150 m^2^/g (Alburquerque et al., 2016).

Based on data from Table 1, at any HC process time and temperature, the C content exceeded 50%, and H/C is lower than 0.7. The surface area (BET) never exceeded 150 m^2^/g, but increased by 120% compared to the original level (89% within 30 min of process time). The O/C ratio started from an original level of 0.44, barely exceeding the threshold, fell to 0.41 after 5 min of process time, later ranging between 0.49 (after 30 min) and 0.88 (highest level after 45 min). While neither BET area nor O/C complied with the quality criteria in the original biochar sample, the HC process led the surface area towards the respective target; on the contrary, the simultaneous decrease of carbon content and the increase of oxygen content led to the observed growth of the O/C ratio.

Fig. 4 shows the O/C evolution throughout the HC process, which deserves a deeper investigation.

**Figure 4.**
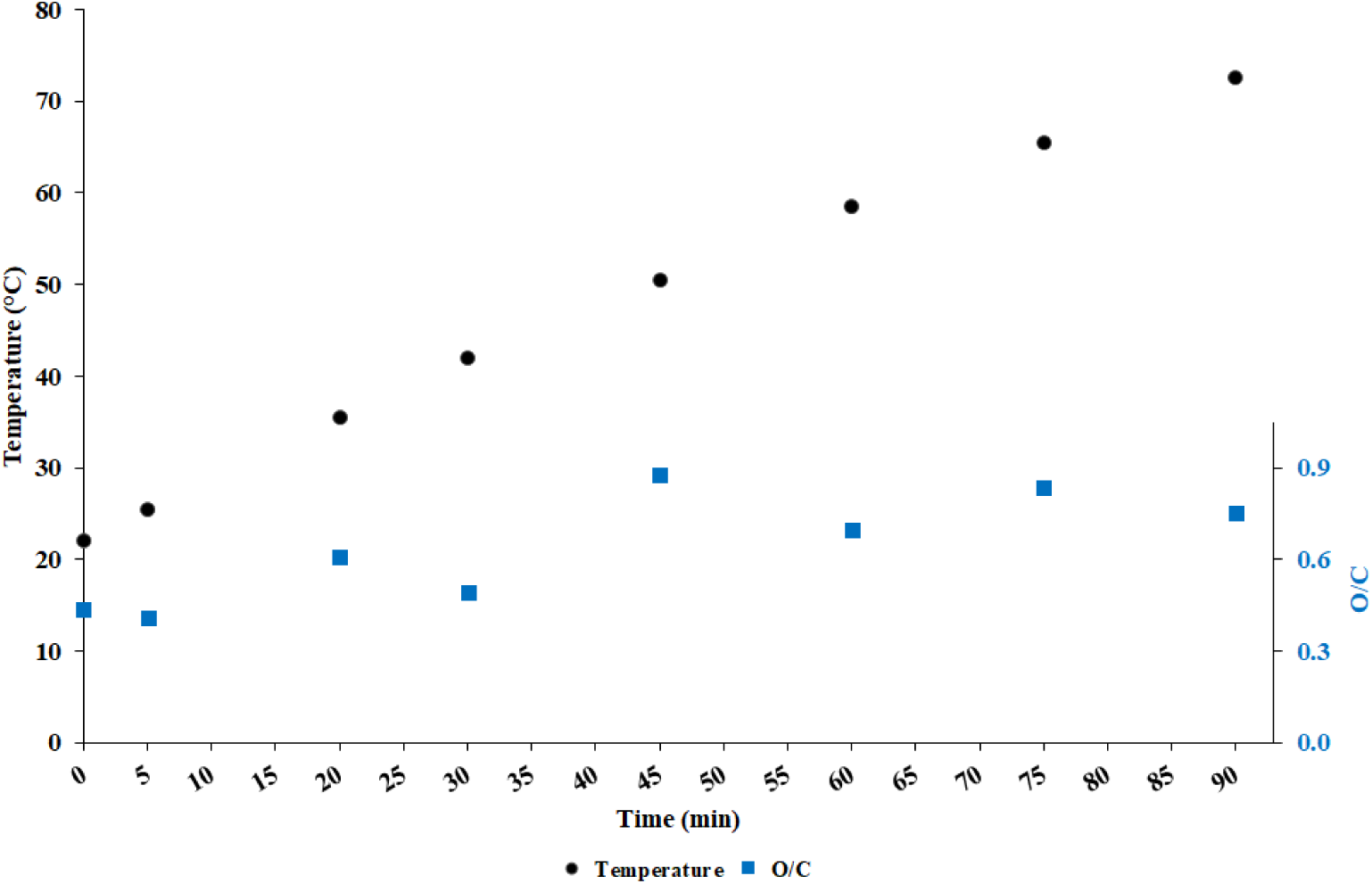
Evolution of the O/C ratio throughout the HC process.

Two oscillating patterns arise, separated by a substantial upward jump from 30 to 45 min, within the temperature range 42 to 50.5 °C, due to the simultaneous decrease of carbon content and increase of oxygen content.

Such jump could be related to the increasing aggressiveness of the cavitation process with increasing temperature, at least up to a certain temperature: in pure water, the peak aggressiveness was observed around 60°C (Dular, 2016), but it could occur at lower temperatures in the study case. In turn, more aggressive cavitation is likely to enhance the generation of molecular oxygen, according to well-known water splitting reactions (Ciriminna et al., 2017), which could be absorbed by the biochar, as well as to enhance carbon solubilization, *i.e.* the release of carbon from the biochar to the aqueous medium (Maeng et al., 2010; Topac Sagban et al., 2017).

The oscillating pattern, in turn, could be explained by some equilibrium kinetics between adsorption and desorption of both carbon and oxygen, but advancing a comprehensive model is outside the scope of this study, as well as including in the picture the process time, *i.e.* the number of passages through the cavitation reactor. However, on the average, the O/C ratio remains close to the original value within the first 30 min of process time, with temperatures lower than 42°C, as well as close to the recommended range. This evidence suggests that the above limits for HC process time and temperature are safe to obtain a useful biochar on the side of the O/C ratio.

Based on data shown in Table 1, as noted in subsection 3.3, the nitrogen content quickly increased after 5 min of process time, to stabilize around 0.58% from 20 min onwards, in comparison to the original level of 0.49%. This consequence of the HC process could be positive, with regards to the following release of that nutrient to agricultural soil and plants (Alburquerque et al., 2016).

The evolution of the ash content shows the respective dropping by more than 20% after only 5 min of process time, before stabilization throughout the process, with a local minimum at the level of 31.21% after 30 min of process time. The pH increased only marginally during the first 30 min of process time (pH 9.35 to pH 9.47), then further increased to pH 9.63 when measured 24 h after the process.

Recalling that the main aim of this study was the increase of the BET surface area by means of the HC process, as an alternative to conventional higher-temperature pyrolysis processes, it can useful to note that, generally, TP leads to substantial increase of the ash content with working temperature. For example, over 60% from 400°C to 600°C using conocarpus wastes (Al-Wabel et al., 2013), or in the range 20% to 80% from 350°C to 550°C using AS, OS, and PW (Gómez et al., 2016), despite a notable exception found in the latter study for biochar based on olive-tree pruning.

Along with ash content, pH was observed to increase substantially with pyrolysis temperature. For example, from pH 9.67 at 400°C to pH 12.21 at 600°C using conocarpus wastes (Al-Wabel et al., 2013), or from pH 10.78 to pH 10.99 using wheat straw, and pH 9.35 to pH 12.95 using lignosulfonate, in slow pyrolysis regime from 400°C to 600°C (Gómez et al., 2016).

The increase of ash content and the excessive alkalization of biochar, due to high pyrolysis temperatures, was shown to be detrimental for soil applications (Mukherjee and Lal, 2014); such issue was avoided with the HC process, especially, with reference to the pH, within 30 min of process time.

Based on the results, and within the limits, of the discussed case study, HC processing with short enough process time, *e.g.* 30 min, and within a moderate temperature range, *e.g.* no more than 42°C, appears a viable and energy efficient way to increase the biochar surface area, in place of increasing the working temperature of conventional pyrolysis processes.

The following main results, for HC processing of biochar generated by a TP process up to a certain working temperature, arise:

- Much higher process yield, an order of magnitude or more, compared with elevating the working temperature in TP processes, the process yield being computed based on energy consumption, biochar yield, BET and carbon content in the biochar;
- Preservation of acceptable levels of carbon concentration, as well as low values of the H/C ratio;
- Practical retention of the original level of the O/C ratio;
- Substantial increase of the nitrogen content;
- Decrease of the ash content, contrary to what happens with increasing working temperature in TP processes;
- Growth of the pH, but limited to levels much lower than observed with TP processes.

## 4. Conclusions

Increasing the porosity and surface area (BET) of biochar is generally desirable due to the respective positive effects on exchange processes in the soil. However, the option to increase the working temperature in conventional TP processes comes with substantial drawbacks, such as increased energy consumption, reduced yield of biochar with respect to the raw material, and an increase of the ash content and the pH, compensated by an increase of the stably sequestered carbon.

The aim of this study was to overcome the above-mentioned drawbacks of TP processes with the aid of HC processes applied to biochar generated at limited TP working temperatures. Even within the limits of this study, HC revealed as a very promising alternative to higher-temperature TP, so much that, despite a marginal decrease of carbon content, but within the acceptable range, HC process yields were at least an order or magnitude greater than for TP processes. Moreover, the original chemical composition of the biochar was preserved, or even improved for the purpose of soil applications.

The results achieved in this study need extensive confirmation. Nevertheless, potentially wide margins for further improvement of HC processing of biochar exist, such as the possibility of avoiding the preliminary sieving or grinding of the biochar by using an open-impeller pump, and increasing the biochar concentration in water, which would further improve the energy balance and the yield of the HC process.

The investigation about the aforementioned potential improvements, as well as, for example, the verification of the effectiveness of HC processing with different types of biochar, or biochar produced from different pyrolysis parameters, or different production processes such as gasification or hydrothermal carbonization, are recommended as subjects for further research.

## Acknowledgements

The authors dedicate this article to the memory of Prof. Giampiero Maracchi, an inspirational leader for all of us.

A special thanks to Dr Massimo Valagussa (MAC - Minoprio Analisi e Certificazioni S.r.l.) and Italian Biochar Association (ICHAR http://www.ichar.org) for supply and characterization of biochar

## Supplementary Material

**Table S1.** Surface Areas and Pore volumes of various biochar before and after cavitation at different times of treatment

| Time<br>(min) | BET-surface<br>area<br>(m<sup>2</sup>/g) | Langmuir<br>surface<br>area<br>(m<sup>2</sup>/g) | *t*-plot-<br>micropore<br>area<br>(m<sup>2</sup>/g) | BJH-desorption-<br>cumulative surface<br>pore area<br>(m<sup>2</sup>/g) | Total pore<br>volume<br>(cm<sup>3</sup>/g) | *t*-plot<br>micropore<br>volume<br>(cm<sup>3</sup>/g) | BJH-Desorption-<br>cumulative pore<br>Volume<br>(cm<sup>3</sup>/g) | BJH average<br>desorption<br>pore width<br>(nm) |
| --- | --- | --- | --- | --- | --- | --- | --- | --- |
| 0 | 59.95 | 101.91 | 35.38 | 10.36 | 0.047 | 0.023 | 0.021 | 7.992 |
| 5 | 80.25 | 136.67 | 47.02 | 17.76 | 0.063 | 0.030 | 0.028 | 6.290 |
| 15 | 95.55 | 162.01 | 60.30 | 15.30 | 0.071 | 0.038 | 0.024 | 6.325 |
| 30 | 113.32 | 192.17 | 71.59 | 21.13 | 0.083 | 0.045 | 0.032 | 6.053 |
| 45 | 130.76 | 222.02 | 86.06 | 27.02 | 0.095 | 0.054 | 0.039 | 5.800 |
| 60 | 135.27 | 229.60 | 89.00 | 27.43 | 0.097 | 0.056 | 0.039 | 5.749 |
| 75 | 129.21 | 219.11 | 86.77 | 26.14 | 0.094 | 0.054 | 0.041 | 6.251 |
| 90 | 131.05 | 222.73 | 86.21 | 26.47 | 0.098 | 0.054 | 0.045 | 6.797 |

## N_2_-adsorption-desorption isotherms and BJH Desorption dV/dLog(w) Pore Volume for Biochar, starting from the sample collected after 45 min of process time

**Fig. S1.**
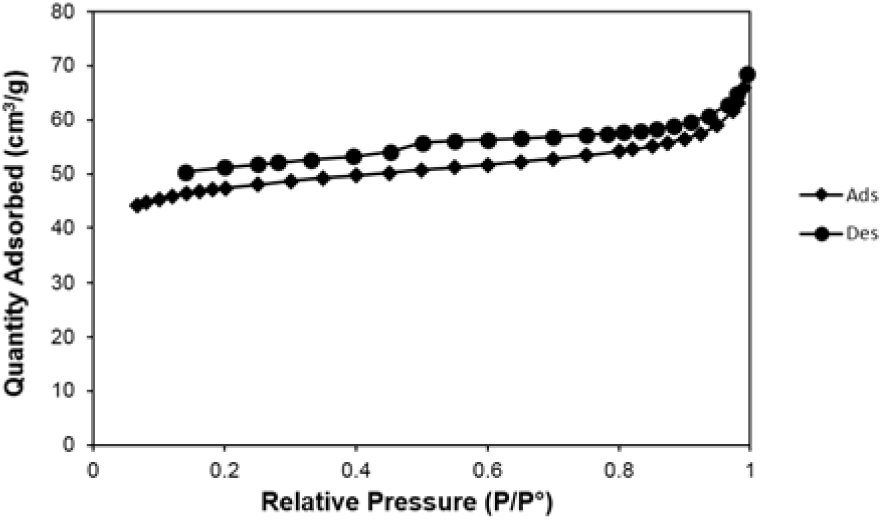
N_2_-adsorption-desorption isotherms at 77.4 K for Biochar after 45 min cavitation treatment

**Fig. S2.**
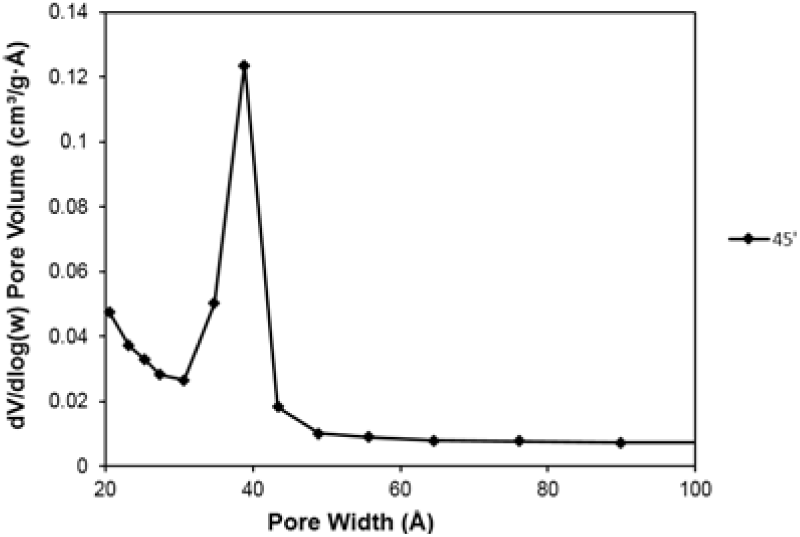
BJH Desorption dV/dLog(w) Pore Volume for Biochar after 45 min treatment

**Fig. S3.**
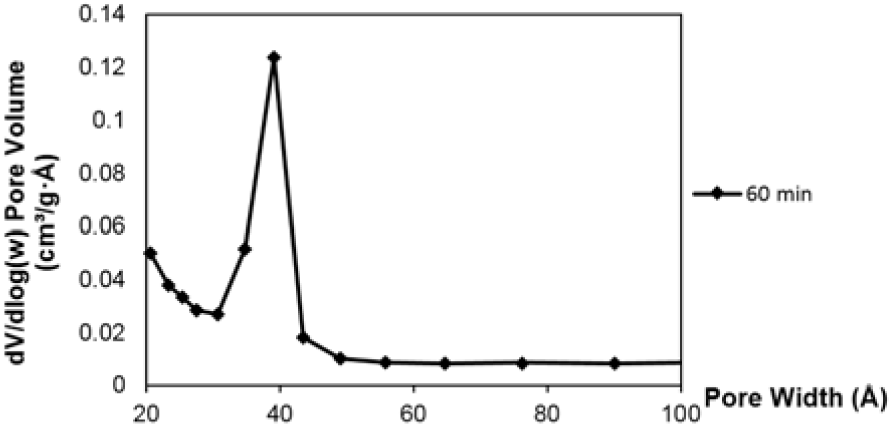
BJH Desorption dV/dLog(w) Pore Volume for Biochar after 60 min treatment

**Fig. S4.**
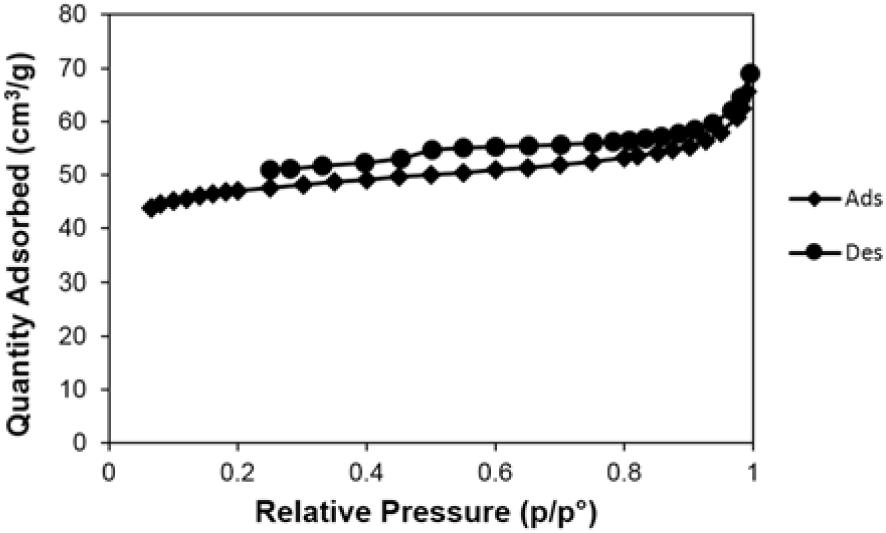
N_2_-adsorption-desorption isotherms at 77.4 K for Biochar after 75 min cavitation treatment

**Fig. S5.**
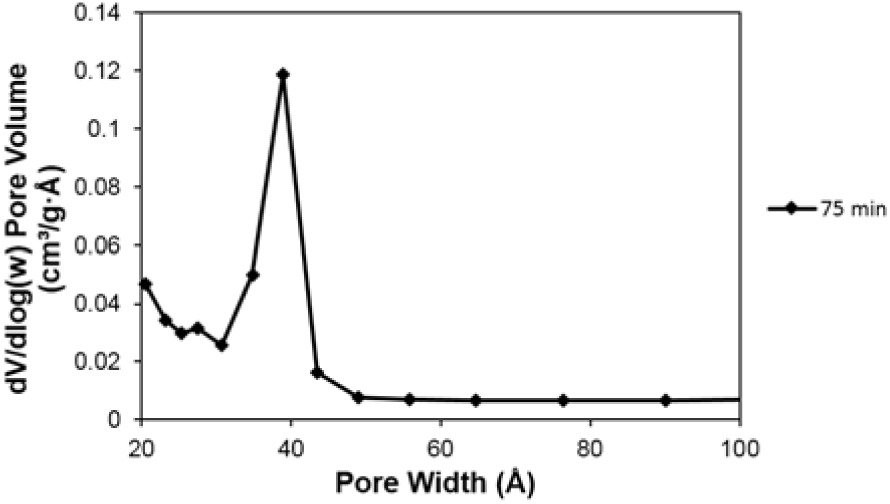
BJH Desorption dV/dLog(w) Pore Volume for Biochar after 75 min treatment

**Fig. S6.**
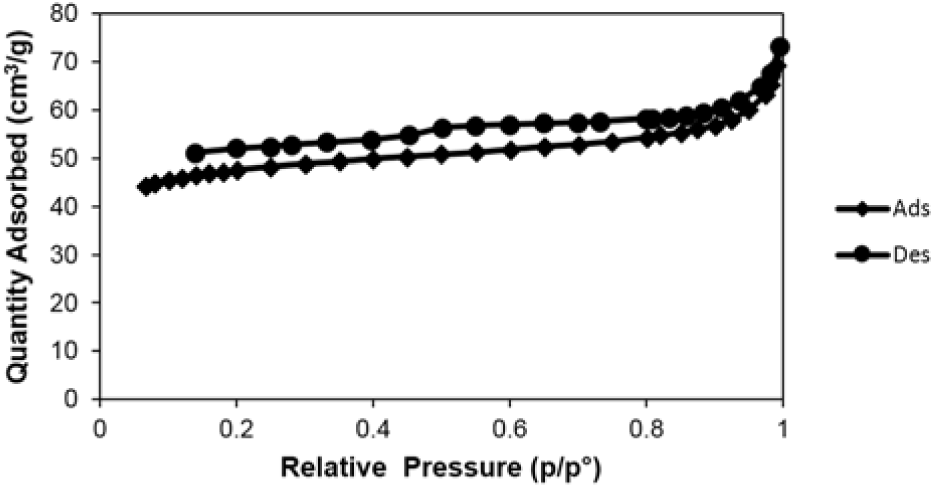
N_2_-adsorption-desorption isotherms at 77.4 K for Biochar after 90 min cavitation treatment

**Fig. S7.**
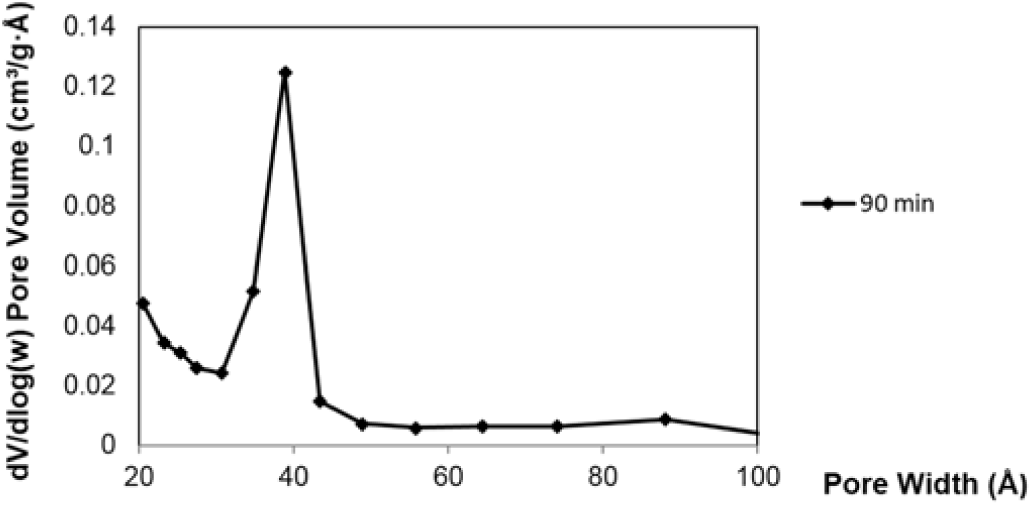
BJH Desorption dV/dLog(w) Pore Volume for Biochar after 90 min treatment

